# Leveraging complex interactions between signaling pathways involved in liver development to robustly improve the maturity and yield of pluripotent stem cell-derived hepatocytes

**DOI:** 10.1101/2020.09.02.280172

**Authors:** Claudia Raggi, Marie-Agnès M’Callum, Quang Toan Pham, Perrine Gaub, Silvia Selleri, Nissan Baratang, Chenicka Lyn Mangahas, Gaël Cagnone, Bruno Reversade, Jean-Sébastien Joyal, Massimiliano Paganelli

**Author notes:** **Correspondence:**, @maxpaganelli.

## Abstract

Pluripotent stem cell (PSC)-derived hepatocyte-like cells (HLC) have shown great potential as an alternative to primary human hepatocytes (PHH) for *in vitro* modeling. Several differentiation protocols have been described to direct PSC towards the hepatic fate, although the resulting HLC have shown more a fetal than adult phenotype. Here, by leveraging recent knowledge of the signaling pathways involved in liver development, we describe a robust, scalable protocol that allows to consistently generate high-quality HLC from both ESC and iPSC. Such HLC are comparable to adult PHH in terms of key mature liver functions and proved suitable to assess drug hepatotoxicity, as a proof of concept of their potential as a physiologically representative alternative for *in vitro* modeling.

## INTRODUCTION

For decades, the study of human liver physiology and development, as well as drug discovery, have been limited by the lack of representative models. Animal studies and *in vitro* models based on tumor cell lines proved crucial to build our current knowledge but are inadequate to assess the specificities of the human liver. Primary human hepatocytes (PHH) suffer from major limitations that have hampered our ability to create proper *in vitro* human models and generate representative data. Although considered the “gold standard” by regulatory agencies, PHH do not allow a reliable prediction of a drug’s metabolism and toxicity in human subjects, which results in a high failure rate at clinical phases and is partially responsible of the high costs and long times of drug development.

Stem cell-derived hepatocyte-like cells (HLC) have shown great potential as an alternative to PHH. Pluripotent stem cells (PSC), such as embryonic stem cells (ESC) and induced pluripotent stem cells (iPSC), are a potentially unlimited source from which to produce HLC.^1^ Various differentiation protocols inspired by liver organogenesis have been described for both ESC and iPSC. Empirical identification of best culture conditions has historically played a central role in establishing such protocols. Our knowledge of liver development has greatly advanced over the last decade and we now have a reasonable understanding of the signaling pathways involved.^2–4^ This helped to improve the quality of PSC-derived HLC.^5–7^ Nevertheless, obtained HLC are still closer to fetal than adult PHH.^8^ Moreover, the quality of differentiation varies widely among HLC obtained from different PSC populations, and cell yield is often low with most of described protocols.^9^ These limitations constitute a significant barrier to the full implementation of HLC for *in vitro* modeling, drug testing and cell therapy applications. By leveraging most recent knowledge of signaling pathways involved in liver development and timely acting on Wnt/β-catenin, oncostatin M (OSM) and TGFβ pathways during the later stages of differentiation, we developed a new robust protocol allowing for high-yield, consistent generation of high-quality bipotent liver progenitors and HLC from both ESC and iPSC.

## RESULTS

We used 6 different PSC populations (one ESC line and five iPSC lines) and differentiated them through consecutive stages following liver development (Fig. 1A,B).

**Figure 1.**
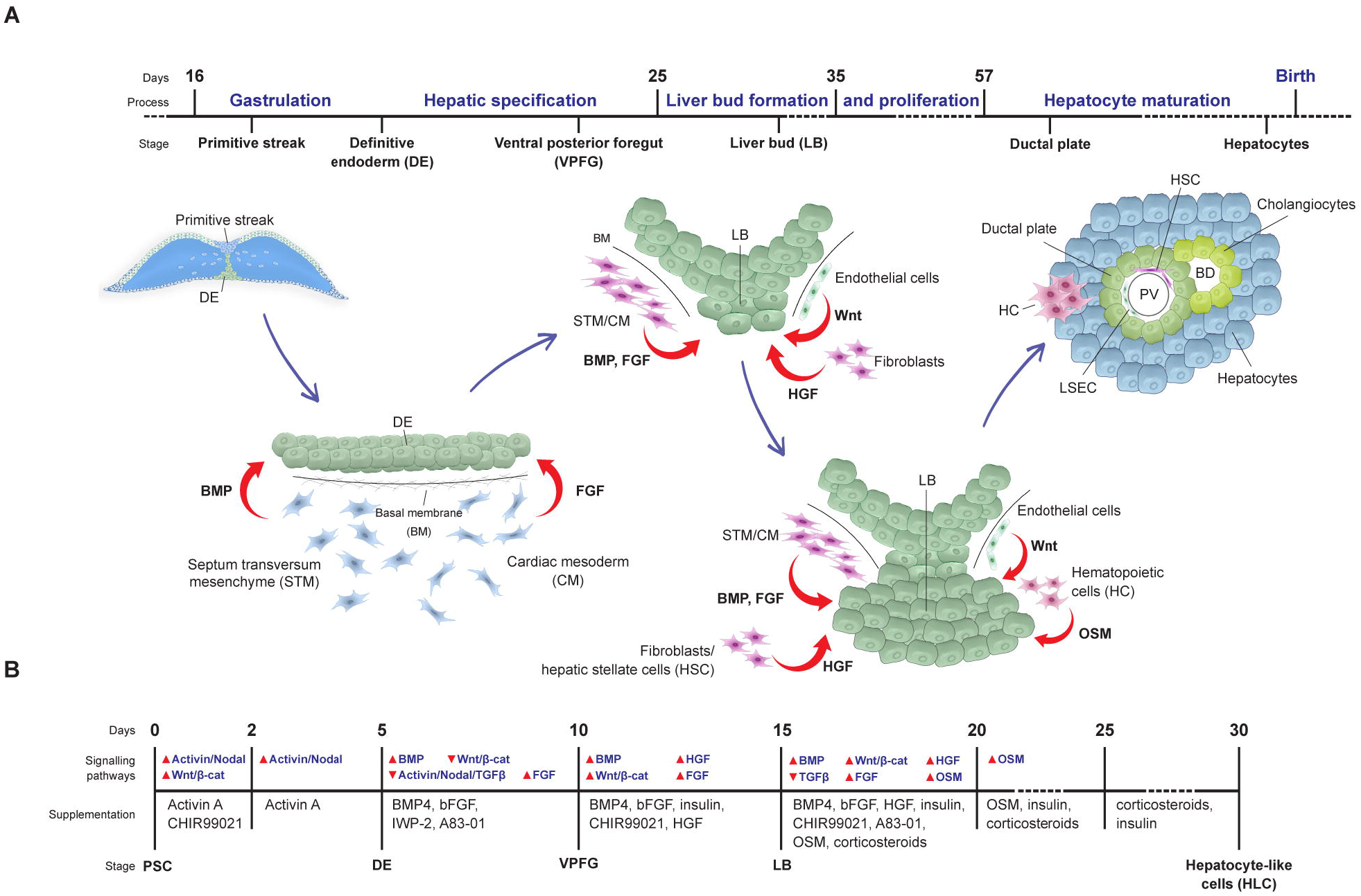
Mimicking liver development. **A)** Schematic representation of human liver development showing the role that the main actors and signaling pathways play at each developmental stage; **B)** Description of our new differentiation protocol showing how the sequential supplementation of activators and inhibitors allows to reproduce the signals acting on the developing liver tissue in the embryo at corresponding stages.

### Hepatic specification

After careful characterization of PSC (Supplementary Fig. S1), we recreated the process of gastrulation *in vitro,* passing by a primitive streak stage and then pushing Brachyury (T)-positive cells towards the mesendoderm through epithelial-mesenchymal transition by concurrently activating both Activin/Nodal signaling (with Activin A) and canonical Wnt/β-catenin pathway (via the selective inhibition of GSK3β with CHIR99021). This led to the timed upregulation of EOMES and FOXA2 (24 hours) and SOX17 (48 hours; Fig. 2A). Stimulation of Activin/Nodal pathway for 3 additional days allowed the cells to acquire the gene and protein expression profile typical of the definitive endoderm (DE; Fig. 2B,C). The obtained DE population was highly homogenous, with a >90% of FOXA2, SOX17 and CXCR4-positive cells (Fig. 2D,E). The addition of knock-out serum replacement (KOSR) at low concentration, which, as fibroblast grow factor (FGF) secreted by the cardiac mesoderm, induces hepatic specification through the MAPK signaling pathway, significantly reduced cell death without affecting the quality of the obtained DE (Supplementary Fig. S2A,B). No difference was seen in the results obtained from applying this protocol to ESC or iPSC populations (Supplementary Fig. S2C).

**Figure 2.**
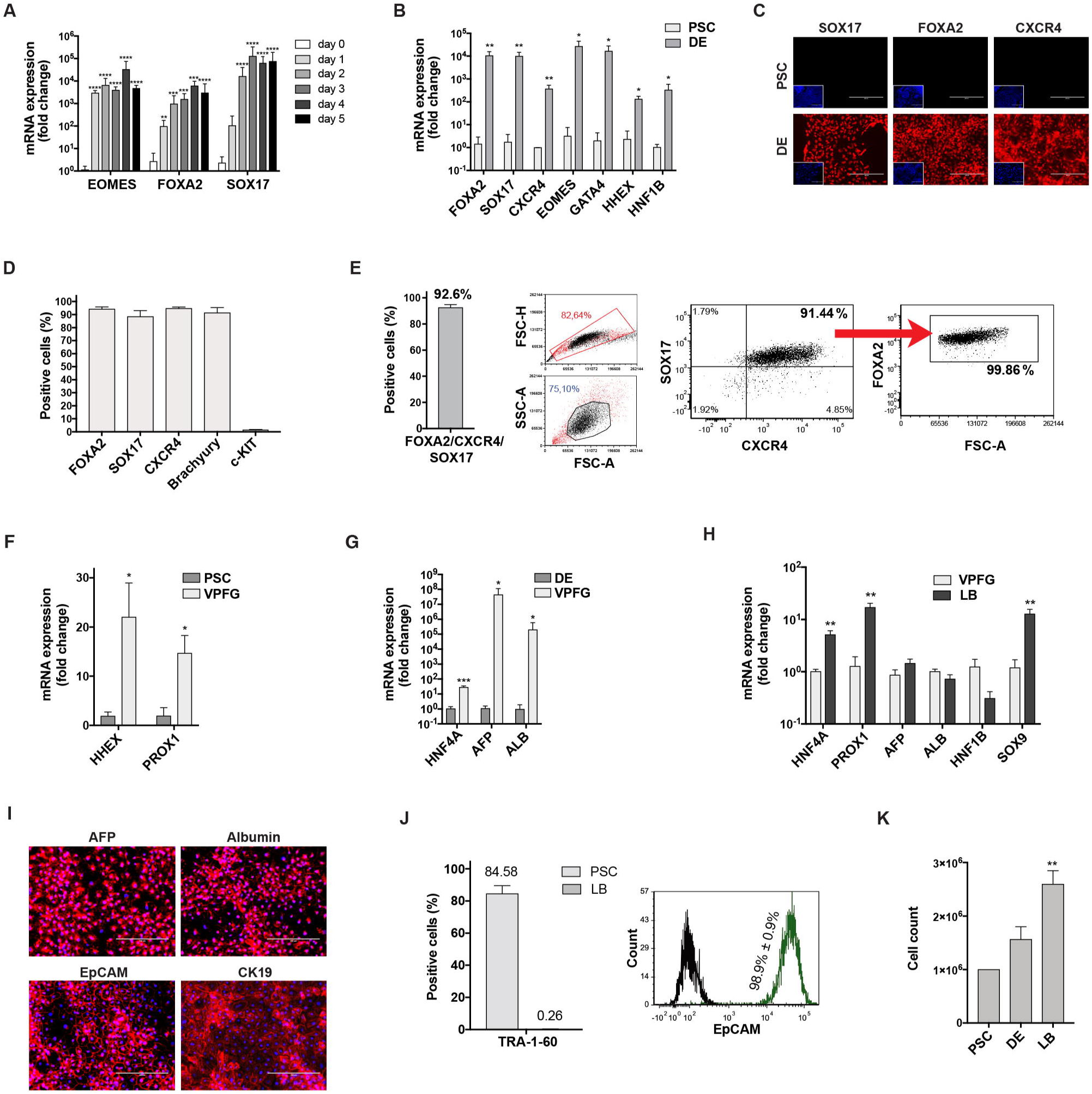
A-E) PSC-derived endoderm (DE). **A)** Expression of EOMES, FOXA2 and SOX17 genes over the first 5 days of differentiation (RT-qPCR, expressed as log_10_ mean fold change ± SEM, n=9, ** p<0.01, *** p<0.001, **** p<0.0001). **B)** Expression of endoderm-enriched genes in PSC-derived cells after 5 days of differentiation compared to undifferentiated PSC (RT-qPCR, expressed as log_10_ mean fold change ± SEM, n=3 for PSC, n=6 for DE, * p<0.05, ** p<0.01). **C)** Expression of DE markers at the end of day 5 of differentiation compared to undifferentiated PSC (immunofluorescence, representative images, DAPI nuclear staining in included images, scale bar 200 μM). **D)** Expression of DE markers is highly homogeneous in PSC-derived DE cells (flow cytometry, n≥4, mean ± SEM). **E)** FOXA2/CXCR4/SOX17 triple-positive cells constitute >90% of PSC-derived DE (flow cytometry, n=4; left: mean ± SEM, center and right: representative experiment). **F-K) Posterior foregut and liver bud formation. F-G)** PSC-derived VPFG: expression of HHEX, PROX1, HNF4A, AFP and ALB genes in PSC-derived cells (RT-qPCR expressed as log_10_ mean fold change ± SEM, compared to: F, undifferentiated PSC and G, PSC-derived DE; n=3 for PSC, n≥3 for DE, n≥5 for VPFG; * p<0.05, *** p<0.001). **H)** PSC-derived cells after 15 days of differentiation achieve a gene expression profile consistent with hepatoblasts in the forming LB (RT-qPCR, expressed as log_10_ mean fold change ± SEM; n=3 for VPFG, n=8 for DE, ** p<0.01). **I)** Expression of hepatoblast-enriched AFP, albumin, EpCAM and CK19 in PSC-derived LB cells (immunofluorescence, representative images with DAPI nuclear staining, scale bar 200 μM). **J)** LB cells show high homogeneity for EpCAM expression, while expression of a marker of pluripotency is negligible and comparable to background noise (flow cytometry; TRA-1-60: n=6 for PSC, n=3 for LB, mean ± SEM; EpCAM: n=5, mean ± SEM, representative experiment). **K)** The differentiation protocol allows the expansion of LB cells (automated cell count, n=6 for PSC and LB, n=3 for DE; mean ± SEM, ** p<0.01).

In the embryo, the DE is then exposed to BMP from septum transversum mesenchyme (STM) and FGF from the cardiac mesoderm (Fig. 1A).^2,3^ This drives the cells towards the formation of the posterior foregut. Inhibition of Wnt signaling pathway is needed to relieve the repression of *HHEX,* which is required for the commitment to the hepatic fate.^10^ TGFβ inhibition allows for upregulation of albumin *(ALB)* and *PROX1* genes, the latter playing an essential role in subsequent liver bud formation.^11^ After 5-day exposure to BMP4, low-dose FGF, A83-01 (TGFβ pathway inhibitor) and IWP2 (Wnt pathway inhibitor), the cells acquired a phenotype suggestive of the commitment of the ventral posterior foregut (VPFG) towards the hepatic fate (corresponding to mouse embryonic day E8.5), with the overexpression of *HHEX, PR0X1* (Fig. 2F), *HNF4a*, α-fetoprotein *(AFP) and ALB* genes (Fig. 2G).

### Liver bud formation and expansion

In humans, 9 days post-gastrulation, hepatic progenitor cells from the VPFG delaminate and invade the surrounding stroma, forming the liver bud (LB). Liver progenitors (hepatoblasts) enter hence into contact with the STM, which continue secreting BMP, and are in close proximity with sinusoidal endothelial cells secreting Wnt and hepatocyte growth factor (HGF; Fig. 1A). Supplementation of hepatoblasts with BMP4, FGF, HGF and CHIR99021, in the presence of insulin, for 5 days, triggered the re-establishment of cell-cell contacts and a deepest commitment towards the hepatic fate (Fig. 1B). At the end of this stage, such LB cells further increased the expression of transcription factors *HNF4a* and of *PR0X1,* both required for their proliferation and differentiation (Fig. 2H).^11,12^*ALB* and *AFP* continued to be very highly expressed in LB cells. The expression of *HNF1b,* which at this stage controls cholangiocyte fate, was instead reduced, while the expression of SRY-box 9 *(SOX9)* was significantly increased in LB cells, which is consistent with the expression of these genes in bipotent hepatic progenitor cells (0Fig. 2H).^13^ Immunostaining showed that 79.1%± 13.4% of LB cells were AFP-positive, 80.4%± 10% were albumin-positive and 97.2%± 2.7% EpCAM-positive (Fig. 2I). The expression of cytokeratin 19 (CK19) was ubiquitous, although variable between cells and significant in only 36.3%± 28% of them. EpCAM staining at the cell surface confirmed the re-establishment of cell-cell contacts (Fig. 2J). The number of cells still expressing pluripotency markers was negligible at this stage (Fig. 2J). The staged activation of HHEX, HNF4a and PROX1 also led to the proliferation of hepatoblasts (Fig. 2K), in line with the behavior of the liver bud at this stage of development (day 24-48 of gestation, corresponding to E10-E14 in mouse).^11^ No difference between ESC and iPSC was noted (Supplementary Fig. S2D).

### Maturation into hepatocyte-like cells

In the developing human embryo, starting at around gestational day 57 (E13.5 in mouse), the close contact between hepatoblasts and hepatic stellate cells (secreting HGF and FGF) and hematopoietic cells (producing OSM) drives their maturation into hepatocytes (Fig. 1A). Mature liver functions are then progressively acquired, with the liver expressing a fully mature phenotype only several weeks after birth. We exposed LB cells to OSM and dexamethasone in addition to the stimulation of BMP, FGF, HGF and Wnt signaling pathways and inhibition of TGFβ pathway, for 5 days, to induce maturation to HLC (Fig. 1B). Subsequently, we maintained only OSM and dexamethasone for another 5 days and then concluded the differentiation processes with further 5 days without OSM, in order to mimic the decrease in the stimulus from hematopoietic cells that happens in the later weeks of gestation.^14^ At the end of the differentiation process, we obtained a homogeneous monolayer of mono- and binucleated polygonal epithelial cells (Fig. 3A). Such PSC-derived HLC showed an important increase in the expression of *ALB* (>1000-fold change) and *AFP* (>10-fold change) compared to LB hepatoblasts (Fig. 3B). The expression of mature liver-specific genes *ALB, ASGR1* and *TAT* in HLC was comparable to PHH (Fig. 3C), with *HNF4a* and *AFP* being more highly expressed. The differentiation protocol proved to be robust, with negligible inter-population and interoperator variability (32 differentiations of 6 PSC populations, 3 different operators; Supplementary Fig. S2E). No difference between ESC and iPSC was noted (Supplementary Fig. S2F). Albumin and AFP expression was very homogeneous in HLC (Fig. 3D). Interestingly, CK19 was expressed by 44.6%± 22.4% of the cells, most of which were also albumin positive. HLC effectively performed mature liver functions such as cytochrome P450 3A4 (Cyp3A4) activity, albumin secretion and urea production and resulted comparable to metabolism-qualified adult PHH (Fig. 3E).

**Figure 3.**
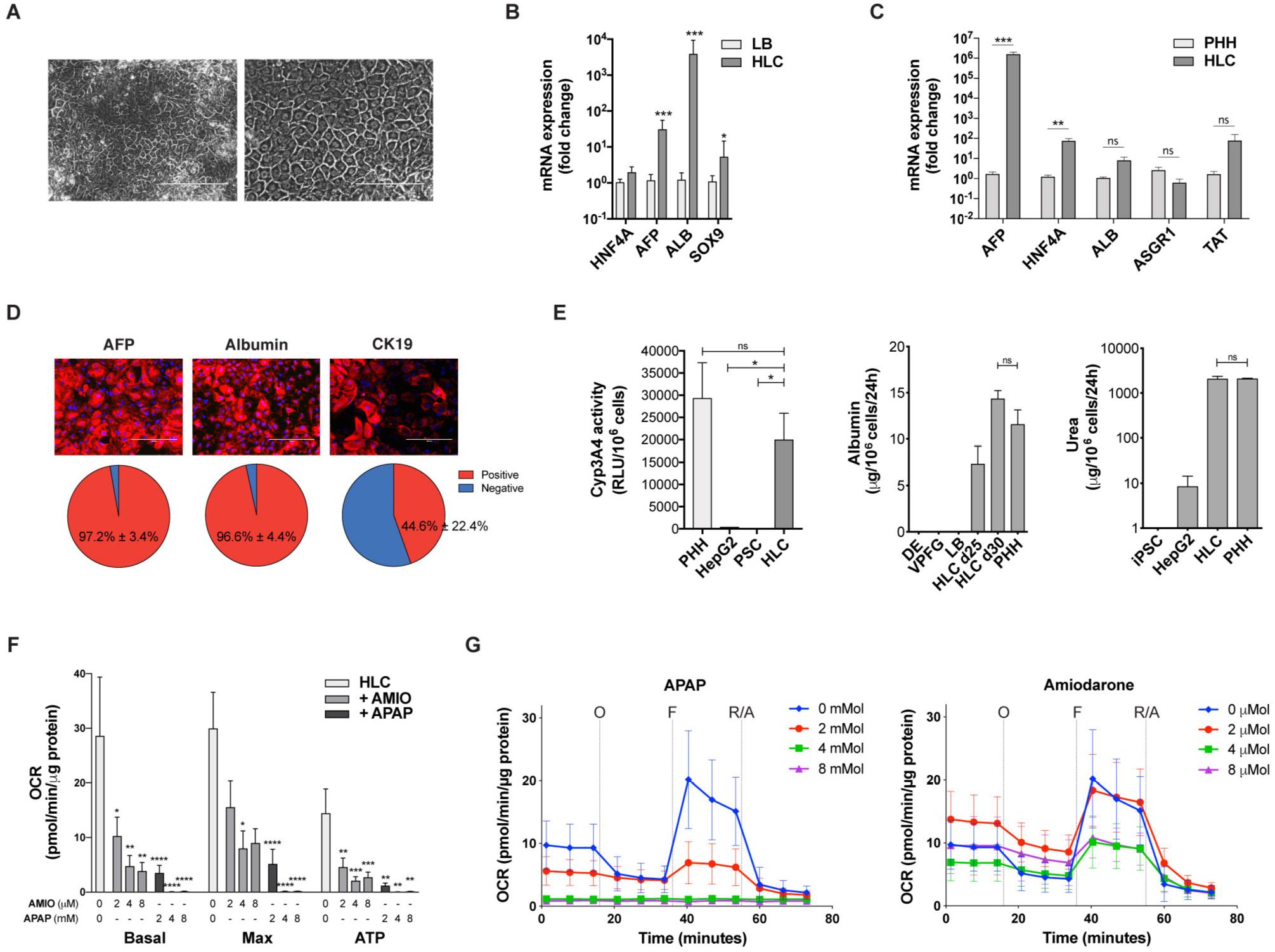
PSC-derived HLC. **A)** Representative morphology at the end of the differentiation protocol (phase contrast, scale bar 200 μM left, 100 μM right); **B)** Expression of liver-specific genes compared to LB cells (RT-qPCR, expressed as log_10_ mean fold change ± SEM; n=7, * p<0.05, *** p<0.001). **C)** Expression of liver-specific genes compared to adult PHH (RT-qPCR, expressed as log_10_ mean fold change ±SEM; n=9 for PHH, n=10 for HLC; ** p<0.01, *** p<0.001, ns p≥0.05). **D)** Expression of liver-enriched AFP, albumin and CK19 in PSC-derived HLC (immunofluorescence; top: representative images with DAPI nuclear staining, scale bar 200 μM; bottom: percentage of positive cells compared to DAPI-positive nuclei, n=9, mean ± SEM). **E)** Liver-specific functions performed by HLC: cyp3A4 activity (expressed as relative light units per million cells and compared to PHH, HepG2 cells and PSC; n=6 HLC, n=7 PHH, n=5 PSC, n=3 HepG2; mean ± SEM, * p<0.05); albumin secretion in HLC at day 25 and 30 compared to previous stages of differentiation and to PHH (ELISA, n=4 for HLC and PHH, n=3 for DE, VPFG and LB, mean ± SEM); urea production (ELISA, n=3 for HLC, PSC and HepG2, n=12 for PHH, mean ± SEM). **F-G)** Hepatotoxicity of acetaminophen (APAP) and amiodarone (AMIO) assessed through the measurement of oxygen consumption rate on HLC (O: oligomycin, F: FCCP, R/A: Rotenone/Antimycin A; n=12, mean ± SEM).

As a proof of concept, we used such HLC to asses hepatotoxicity of two common drugs with a known dose-dependent effect on hepatocytes (Fig. 3F,G). Mitochondrial dysfunction has emerged as an important mechanism and indicator of drug-induced hepatotoxicity.^15^ By measuring their mitochondrial respiration (oxygen consumption rate, OCR), we showed that HLC allow to representatively assess, with very little variability, the toxic effect of acetaminophen and amiodarone on hepatocytes’ metabolism.

When compared to previously published protocols for which a transcriptome was available ^7,16–18^ and to a differentiation protocol not acting on the Wnt/β-catenin and TGFβ signaling pathways beyond the definitive endoderm stage (Supplementary Fig. S3), our new protocol resulted in significantly more mature HLC (Fig. 4 and Supplementary Fig. S4). HLC obtained with our protocol significantly overexpressed key liver-specific genes compared to other previously described HLC and resulted in expression profiles more similar to PHH (Fig. 4A-C). A three-fold increase was found for those encoding albumin and enzymes involved in xenobiotic metabolism, iron homeostasis, and lipids and bile acid metabolism (Supplementary Fig. S4B,C). Analysis of Gene Ontology terms confirmed the overexpression of superclusters of genes that are essential for the development, structural integrity and function of the human liver (Supplementary Fig. S4D). The advantage of the timed regulation of Wnt/β-catenin and TGFβ signaling pathways was even more evident when measuring mature liver functions (Fig. 4E). Moreover, our new protocol allowed to obtain significantly more HLC (Fig. 4F) and resulted in an upregulation of Hippo/YAP and Notch signaling pathways, which is suggestive of the regenerative/replicative state of the cells (Supplementary Fig. S4D) and well explains the better yield.^19^

**Figure 4.**
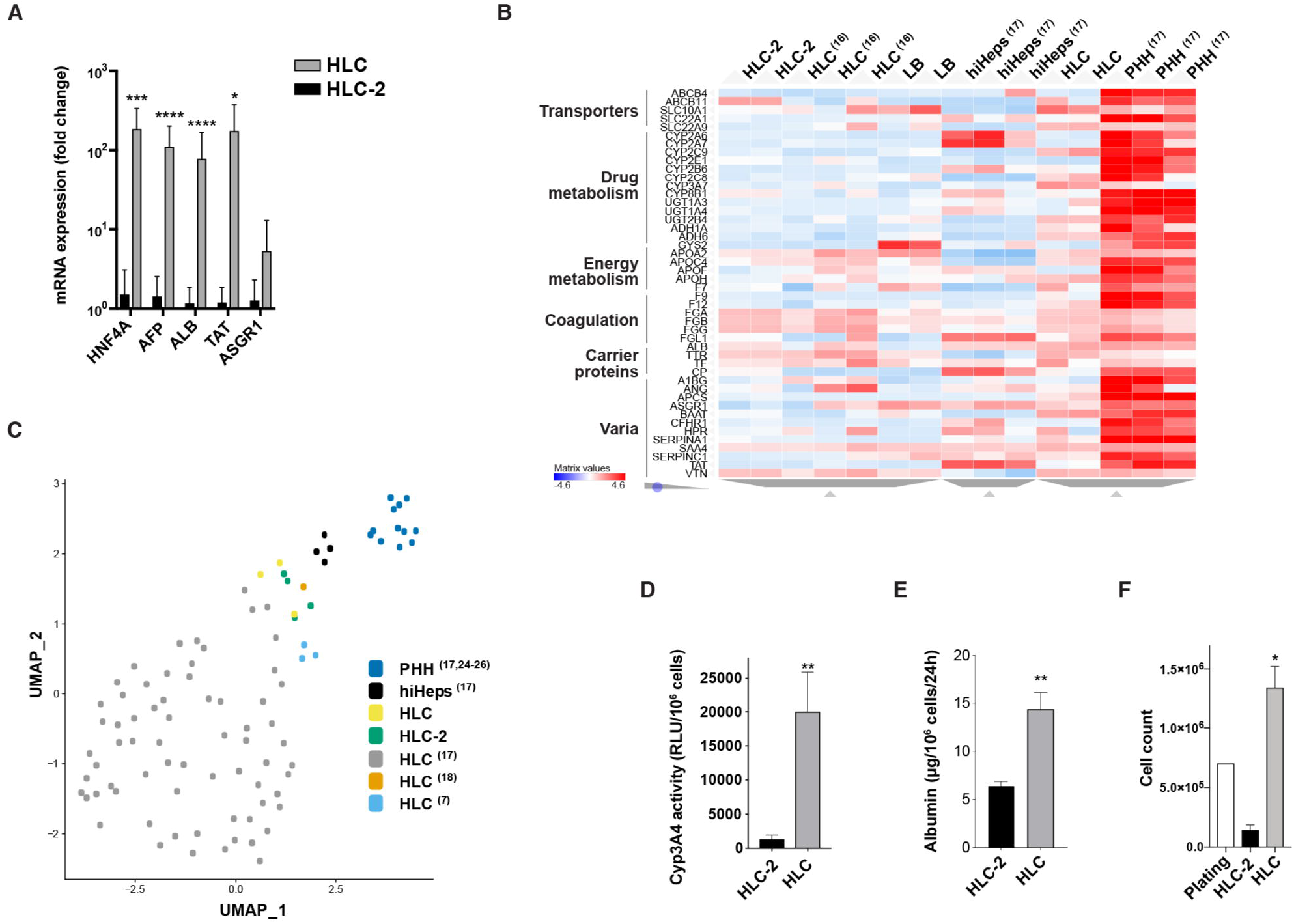
The new differentiation protocol allows obtaining more mature HLC in greater number. **A)** Liver-specific genes are overexpressed in HLC obtained with our new protocol compared to a protocol not acting on Wnt and TGFβ signaling pathways beyond the DE stage (HLC-2; RT-qPCR, expressed as log_10_ mean fold change ± SEM, n=9, * p<0.05, *** p<0.001, **** p<0.0001). **B)** Heatmap showing the expression of the top 46 liver-enriched genes in HLC obtained with our protocol compared to LB cells, HLC-2, HLC obtained with other previously described protocols, hepatocytes derived by transdifferentiation (hiHeps) and PHH (RNA-Seq, unsupervised clustering; list of genes from the Human Protein Atlas).^16,17,23^ **C)** Our HLC are more similar to PHH than most HLC obtained with previously described protocols: 2D representation of Uniform Manifold Approximation and Projection (UMAP)-based dimensionality reduction of the top 17 principal components obtained analyzing the 3000 most variable genes across datasets.^7,16–18,24–27^ **D)** HLC obtained with our new protocol show significantly more Cyp3A4 activity than HLC-2 (relative light units per million cells; n=6 HLC, n=4 HLC-2; mean ± SEM, ** p<0.01). **E)** Our HLC secrete more albumin per million cells than HLC-2 (ELISA, n=4 for HLC, n=10 for HLC-2; mean± SEM, ** p<0.01). **F)** The new differentiation protocol allows generating HLC with significantly more efficiency (automated cell count, n=4; mean ± SEM, * p<0.05).

Overall, our new differentiation protocol allowed to consistently generate more mature HLC from both ESC and iPSC compared to previously described approaches, with good yield and using reagents that have Good Manufacturing Practices (GMP)-compliant versions available. Designed for robustness and ease of clinical translation, the protocol has been licensed to a regenerative medicine company, validated by 3 more independent operators and successfully tech-transferred to an independent contract development and manufacturing organization (CDMO) for follow-on scale-up and optimization.

## DISCUSSION

Over the years, the differentiation protocols were progressively refined to generate more mature HLC from either ESC or iPSC by acting on very diverse signaling pathways. Nevertheless, despite the heterogeneity of growth factors, small molecules, culture media and coating compounds used, full maturity of adult hepatocytes has never been achieved *in vitro,* and a significant variability in the performance of many protocols across various pluripotent stem cell lines was often observed.^8,9^ Here we described a new protocol that, through the timed activation and inhibition of key signaling pathways playing a role along liver development, allows to reproducibly generate high-quality HLC from PSC. This protocol generates highly homogeneous populations at each stage of hepatogenesis, with no significant variability when applied to various iPSC and ESC populations. The obtained HLC were more mature than cells generated with previously described, widely used protocols not leveraging the timed and complex interaction between signaling pathways. Wnt/β-catenin signaling plays a crucial role in orchestrating liver development.^2–4^ Although the role of this pathway during the later stages of development has been well shown in rodents, the need for its subsequent inhibition and activation in synergy with BMP, FGF, OSM and TGFβ signaling modulation has not been previously exploited to generate HLC from PSC. Our HLC expressed most of liver-specific genes responsible for the organ’s synthetic and metabolic activity and performed significantly better at key mature liver functions than previously described protocols not simultaneously acting on such signaling pathways. Only hepatocytes obtained from fibroblasts by transdifferentiation through forced expression of specific hepatocyte transcription factors (hiHeps) showed a gene expression profile more similar to PHH, although they clustered further than our HLC when only key liver-specific genes were considered.^17^ Most importantly, our HLC showed Cyp3A4 activity and albumin and urea production capabilities comparable to adult PHH, and proved suitable for *in vitro* drug testing. Although metabolism-qualified cryopreserved PHH are known to lose functionality compared to freshly isolated PHH, they are still considered the industry “gold standard” to assess candidate new drugs, despite the high inter-donor variability and high costs.^20^

It is widespread experience that most differentiation protocols have an overall low yield, with significant cell death, which increases the costs of differentiation and limits scalability. The activation of Wnt/β-catenin pathway was shown to have an important role on hepatocyte proliferation and might contribute to the good yield of our protocol.^21^ Such a better yield compared to protocols not relying as much on the modulation of TGFβ and Wnt pathways can also be explained by the upregulation of Hippo/YAP signaling shown above. Besides promoting cell proliferation, the activation of this pathway reduces apoptosis, and seem to play a key role in liver regeneration.^19,22^ Sustained Wnt activation during maturation of hepatoblasts into HLC and timed supplementation of OSM to mimic the physiological role of sinusoidal and hematopoietic cells residing in the liver during fetal life allowed for better maturation of our HLC compared to the ESC-derived cells previously described by Touboul et al.^7^ Avoidance of *NOTCH* inhibition resulted in higher *SOX9* expression, but had no impact on mature liver function, while it further supported cell proliferation and reduced the cost of differentiation. Moreover, we avoided the use of serum and used reagents that are easily transitioned to GMP-compliant versions. This will reduce times and costs of GMP implementation and further eases the protocol’s translation for future potential therapeutic applications. Successfully tech-transferred to a cell therapy company and a CDMO, the protocol confirmed its robustness, although optimization will be needed to allow large scale production required for meaningful *in vitro* drug testing and cell therapy applications.

Overall, we described here a new differentiation protocol to consistently obtain high-quality, homogeneous and functional HLC with a high yield from both human ESC and iPSC. With low variability between starting PSC lines, this protocol might prove useful for liver disease modeling. Moreover, the protocol design and yield suggest good potential for scalability, which makes it promising for *in vitro* drug testing and development. Once their potential to replace mature liver functions *in vivo* proven, such HLC might also be considered for cell therapy applications.

## EXPERIMENTAL PROCEDURES

A detailed description of all experimental procedures is provided as Supplementary material.

### PSC differentiation into hepatocyte-like cells (HLCs)

PSCs grown in Essential 8 Flex medium (see Supplementary material for details on PSC generation, characterization and maintenance) were dissociated by TrypLE (Life Technologies) to single cells and seeded on human recombinant laminin 521 (BioLamina)-coated plates in Essential 8 Flex medium at a density of 7 x 10^4^ cells/cm^2^. Differentiation was started (day 0) when the cells reached 70% confluence by changing the medium to RPMI-B27 minus insulin (Life Technologies) supplemented with 1% KOSR (Life Technologies). For the first 2 days, the cells were exposed to 100 ng/ml Activin A (R&D Systems) and 3 μM CHIR-99021 (Stem Cell Technologies), and then for the 3 following days to 100 ng/ml Activin A alone. Subsequently, RPMI-B27 (minus insulin) medium was supplemented with 20 ng/ml BMP4 (Peprotech), 5ng/ml bFGF (Peprotech), 4 μM IWP-2 (Tocris), and 1 μM A83-01 (Tocris) for 5 days, with daily medium change. At day 10, the medium was changed to RPMI-B27 (Life Technologies), supplemented with 2% KOSR, 20 ng/ml BMP4, 5 ng/ml bFGF, 20 ng/ml HGF (Peprotech) and 3 μM CHIR-99021 for 5 days, with daily medium change. At day 16, the medium was changed to HBM/HCM (minus EGF) medium (Lonza), supplemented with 1% KOSR, 20 ng/ml HGF, 20 ng/ml BMP4, 5 ng/ml bFGF, 3 μM CHIR-99021, 10 μM dexamethasone (Sigma), and 20 ng/ml OSM (R&D System) for 5 days, with daily medium change. From day 20, for 5 days, HBM/HCM medium was supplemented with 1% KOSR, 10 μM dexamethasone and 20 ng/ml OSM, changing the medium every other day. From day 25, the cells were maintained in HBM/HCM 1% KOSR medium supplemented with 10 μM dexamethasone, with every other day medium change. During all the differentiation process, the cells were kept at 37°C, ambient O_2_ and 5% CO_2_. Cells were characterized by real-time RT-qPCR, RNA-seq, flow cytometry and immunofluorescence (see Supplementary material for details).

### Functional assessment of HLC

Albumin production was evaluated using the Multigent^®^ microalbumin assay on the Architect cSystems. Cyp3A4 activity and urea synthesis were measured using P450-Glo™ Assay (Promega) and Quantichrom urea assay kit (Gentaur), respectively, according to manufacturers’ instructions.

### Mitochondrial respiration analysis

Mitochondrial stress testing was carried out using a Seahorse Bioscience XF96 analyzer (Seahorse Bioscience) in 96-well plates at 37°C as per the manufacturer’s instructions, with minor modifications (see Supplementary material for details).

### Statistical analysis

Replicates refer to independent experiments on ≥3 different PSC populations. Values are shown as mean ± standard error (SEM). Mann-Whitney U test was used to compare RT-qPCR and functional data. A p-value <0.05 was considered significant.

## ACCESSION NUMBERS

The RNA sequencing data discussed in this publication have been deposited in the NCBI Gene Expression Omnibus under accession number GEO: GSE152390.

## ACKNOWLEDGEMENTS

This work was supported by: Stem Cell Network grant FY17/DT8 (M.P.), Canadian Institute of Health Research New Investigator in Maternal, Reproductive, Child & Youth Health grant MY6-155373 (M.P.), “Fonds de Recherche du Québec – Santé” Junior 1 Clinician-Scientist grant (M.P.). We thank the Vision Health Research Network for funding the Single-cell Academy platform and Ines Boufaied (CHU Sainte-Justine’s Flow Cytometry core facility) for her precious collaboration.

## AUTHOR CONTRIBUTIONS

Conception and design: C.R., M.P.; collection of data: C.R., MA.MC, Q.T.P., P.G., S.S., N.B., CL.M.; data analysis and interpretation: C.R., Q.T.P., G.C., JS.J., M.P.; manuscript writing: C.R., JS.J., M.P.; provision of study material: B.R.; financial support and supervision: M.P. All authors approved the final version of the manuscript.

