## Supplementary material for "Leveraging complex interactions between signaling pathways involved in liver development to robustly improve the maturity and yield of pluripotent stem cell-derived hepatocytes"

### DETAILED MATERIALS AND METHODS

**Ethical approval**

This project was approved by the CHU Sainte-Justine internal review board (CER#1386 and CER#2126) and by the Canadian Institute of Health Research’s Stem Cell Oversight Committee.

**PSC origin, generation and maintenance**

Human ESC (H9/WA09 hESC) were obtained from WiCell Research Institute (Madison, WI, Wisconsin, USA). Human iPSC lines AU24 and HAN3 were provided by Dr. Bruno Reversade. Human iPSC lines FBAT002, HT1A, HT1B were reprogrammed from skin fibroblasts and PBMCs of donors by Sendai-virus delivery using the CytoTune-iPS 2.0 Sendai Reprogramming Kit (ThermoFisher Scientific), following the manufacturer’s instructions. The colonies with suggestive morphology were manually picked based on live staining for the pluripotency marker TRA-1-60 and then cultured as iPSC thereafter, with single colony sub-cloning for the first 5 passages. After passage 10, a temperature shift (incubation at 39°C for 72h) was performed to remove the cMyc gene, and the absence of Sendai virus was confirmed by RT-qPCR. PSC cultures were maintained on vitronectin-coated plates (ThermoFisher Scientific) in Essential 8 Flex medium (ThermoFischer Scientific) at 37°C in humidified atmosphere containing 5% CO_2_ and 4% O_2_.

**PSC differentiation into hepatocyte-like cells (HLCs)**

PSCs grown in Essential 8 Flex medium were dissociated by TrypLE (Life Technologies) to single cells and seeded on human recombinant laminin 521 (BioLamina)-coated plates in Essential 8 Flex medium at a density of 7 x 10^4^ cells/cm^2^. Differentiation was started (day 0) when the cells reached around 70% confluence by changing the medium to RPMI-B27 minus insulin (Life Technologies) supplemented with 1% knockout serum replacement (KOSR, Life Technologies). For the first 2 days, the cells were exposed to 100 ng/ml Activin A (R&D Systems) and 3 μM CHIR-99021 (Stem Cell Technologies), and then for the 3 following days to 100 ng/ml Activin A alone. Subsequently, RPMI-B27 (minus insulin) medium was supplemented with 20 ng/ml BMP4 (Peprotech), 5ng/ml bFGF (Peprotech), 4 μM IWP-2 (Tocris), and 1 μM A83-01 (Tocris) for 5 days, with daily medium change. At day 10, the medium was changed to RPMI-B27 (Life Technologies), supplemented with 2% KOSR, 20 ng/ml BMP4, 5 ng/ml bFGF, 20 ng/ml HGF (Peprotech) and 3 μM CHIR-99021 for 5 days, with daily medium change. At day 16, the medium was changed to HBM/HCM (minus EGF) medium (Lonza), supplemented with 1% KOSR, 20 ng/ml HGF, 20 ng/ml BMP4, 5 ng/ml bFGF, 3 μM CHIR-99021, 10 μM dexamethasone (Sigma), and 20 ng/ml Oncostatin M (OSM, R&D System) for 5 days, with daily medium change. From day 20, for 5 days, HBM/HCM medium was supplemented with 1% KOSR, 10 μM dexamethasone and 20 ng/ml OSM, changing the medium every other day. From day 25, the cells were maintained in HBM/HCM 1% KOSR medium supplemented with 10 μM dexamethasone, with every other day medium change. During all the differentiation process, the cells were kept at 37°C, ambient O_2_ and 5% CO_2_.

**Alternative differentiation protocol not acting on Wnt and TGFβ signaling pathways beyond the definitive endoderm stage**

For comparison, PSCs were differentiated into hepatocyte-like cells (HLC) using a protocol not acting on Wnt and TGFβ signaling pathways beyond the definitive endoderm stage (Suppl. Fig. 3). This protocol was inspired by what previously described by Si-Tayeb et al. with significant modifications.^1^ Briefly, iPSC grown in Essential 8 Flex medium kit were dissociated using TrypLE to single cells and seeded on human recombinant laminin 521-coated plates. The differentiation was started when the cells reached 70-80% confluence. Day 1 and 2 media was changed to RPMI/B27 minus insulin 1% KOSR, supplemented with 100 ng/ml Activin A, 20 ng/ml bFGF, 3 μM CHIR-99021. The addition of KOSR was necessary to limit cell death along the entire differentiation process. From the next 3 days cells were exposed just to 100 ng/ml Activin A. From day 6 for 5 days, media was changed to RMPI/B27 1% KOSR supplemented with 20 ng/ml BMP4 and 10 ng/ml bFGF with daily medium change. From day 11 for 5 days media was changed to RPMI/B27 1% KOSR supplemented with 20 ng/ml HGF. From day 16 to day 28 media was changed to HBM/HCM (minus EGF) supplemented with 20 ng/ml OSM and 10 μM dexamethasone, with every other day medium change. The differentiation was carried out at 37°C, ambient O_2_, and 5% CO_2_.

**Primary human hepatocytes**

Metabolism-qualified primary human hepatocytes (PHH) were obtained from ThermoFisher Scientific (lots Hu1797, Hu1601, Hu8272, Hu2001 and Hu8295), seeded on collagen type 1-coated plates (Corning) and maintained in HBM/HCM medium (Lonza). Analysis were conducted between day 2 and day 6 after seeding.

**Real-time RT-PCR**

Total RNA was extracted using ReliaPrep RNA Miniprep system (Promega) from cultured cells and used as a template for synthesis of single-strained cDNA. Reverse transcription was performed to obtain cDNA (Omniscript, Qiagen). The PCR reaction mixes were prepared and afterwards loaded in the plates. The plates were sealed, centrifuged and then loaded in the instrument (Life Technologies). Standard TaqMan qPCR reaction conditions were used. TaqMan gene expression assays (Thermo Fisher) used are listed in Supplementary table 1. Data were analyzed using the comparative CT (ΔΔCT) method for calculating relative quantitation of gene expression.

**Flow cytometry**

Cells were harvested and aliquoted (0.5 to 1 x 10^6^ cells in each assay tube). They were stained with 100 μl of fluorochrome-conjugated primary antibody solution (membrane antigens) for 20 minutes at room temperature protected from light. Cells were subsequently fixed with 4% paraformaldehyde for 10 minutes at room temperature. In case of intracellular staining, cells were permeabilized with 1% Triton X-100 after washing, and then stained with 100 μl of fluorochrome-conjugated primary antibody solution for 20 minutes at room temperature, protected from light. After extensive washing, cells were resuspended in 0.5 ml staining buffer (1% PBS-BSA) and kept at 4°C until analysis. The used fluorochrome-conjugated primary antibodies were: Per-CP-Cy 5.5 anti-human SOX17 (BD Bioscience), APC anti-human CD184 (CXCR4) (BD Bioscience), PE anti-human FOXA2 (BD Bioscience), PE anti-human EpCAM (BD Bioscience), FITC anti-human TRA1-60 (BD Bioscience), Alexa 647 anti-human Nanog (BD Bioscience), APC anti-human Brachyury (Bio-Techne), PerCP-Cy 5.5 anti-human c-Kit (CD117) (BD Bioscience), PE anti-human SSEA4 (Stem Cell Technologies) and PerCP-Cy 5.5 anti-human TRA-1-81 (BD Bioscience).

**Immunofluorescence on cells**

Cells were fixed in 4% paraformaldehyde for 15 minutes at room temperature. They were subsequently washed with PBS and then permeabilized in 0.2% Triton X-100 for 5 minutes at room temperature. Nonspecific sites were blocked incubating the cells with a 3% blocking serum solution (chosen according to the primary antibody used) for 30 minutes at room temperature. Cells were incubated with primary antibody solution (antibodies were diluted in 2% PBS-BSA) for 1h at room temperature. After washing with PBS, cells were incubated with secondary labelled antibody solution (Alexa Fluor, Life Technologies) for 30 minutes at room temperature, protected from the light. During the last 15 minutes, a dye (Pureblue nuclei staining, BioRad) was added to stain the nuclei. After washing with PBS, cells were mounted with an antifade reagent (ProLong Gold, Life Technologies). Fluorescence was analyzed the day after the procedure using an EVOS FL II microscope (Life Technologies). Used primary antibodies are listed in Supplementary table 2.

**Functional assessment of HLC**

Albumin production was evaluated using the Multigent® microalbumin assay, a quantitative measurement of albumin on the Architect *c*Systems, a turbidimetric immunoassay that uses polyclonal antibodies against human albumin. When the specimen is mixed with the reagents, albumin in the specimen combines with the anti-human albumin antibody (goat) in the reagent to yield an insoluble aggregate that causes increased turbidity in the solution. The degree of turbidity is proportional to the concentration of albumin in the specimen and can be measured optically. Reagent kit: 2K98-20 MULTIGENT Microalbumin. Cyp3A4 activity was evaluated using “P450-Glo™ Assays” from Promega, according to manufacturer’s instructions. Urea synthesis was measured using “Quantichrom urea assay kit” from Gentaur, according to manufacturer’s instructions.

**Mitochondrial respiration analysis**

Mitochondrial stress testing was carried out using a Seahorse Bioscience XF96 analyser (Seahorse Bioscience Inc.) in 96-well plates at 37 °C as per the manufacturer' s instructions with minor modifications. Briefly, cells were seeded at 1x10^5^ cells/well and pre-treated with different doses of acetaminophen (2, 4, 8mM) and amiodarone (2, 4, 8, 19 μM) 24h prior to the assay. On the test day, the growth media was removed, washed twice and replaced with XF assay media (unbuffered DMEM, d5030 Sigma, 25mM Glucose, 2mM Glutamine, 1mM Sodium Pyruvate, pH7.4) and the plate was incubated in a C0_2_-free incubator for 1 h at 37 °C. The hydrated cartridge sensor was loaded with the appropriate volume of mitochondrial modulators to achieve final concentrations in each well: oligomycin (2 µM), carbonilcyanide p-triflouromethoxyphenylhydrazone (FCCP) (2 µM) and with rotenone/antimycin A (both 1 µM). Then, levels of basal respiration, ATP production, proton leak, maximal respiration and non-mitochondrial respiration were analyzed from the OCR values as described in manufacturer's protocol.

**Statistical analysis**

Replicates refer to independent experiments on ≥3 different PSC populations. Values are shown as mean ± standard error. Mann-Whitney U test was used to compare qRT-PCR and functional data. A p-value of <0.05 was considered significant.

**RNA-seq analysis**

Total RNA was isolated as described above. The RNA was then quantified (NanoDrop 8000, ThermoFisher Scientific), and quality controlled (BioAnalyzer 2100, Agilent). 500 ng of total RNA was used as input for the Illumina TruSeq RNA v2 mRNA Sample Prep Kit (Illumina) and sequencing libraries were generated according to the manufacturer’s protocol. The libraries were then pooled and sequenced on a HiSeq 2500 (Illumina) instrument as per the manufacturer’s instructions, generating single-end reads with a length of 100 bp. Following sequencing, fastq files were annotated using Salmon’s quasi-mapping and Ensebl EnsDb.Hsapiens.v86 annotation package.^2^ Quantification of gene expression was performed with Salmon.^2^ The obtained gene counts were then analyzed using the DESeq2 package.^3^ Heatmaps were obtained using heatmap.2 after elimination of lowly expressed genes (cpm > 0.7) and normalization. The lists of liver-enriched genes used for heatmaps were obtained from the Human Protein Atlas (http://www.proteinatlas.org).^4^ Prior to differential expression analysis, genes with low reads maximum read count across samples (<10) were removed. The function DESeq was used for differential expression analysis with default settings. Differentially expressed genes with an adjusted p-value of <0.05 and an absolute log2 fold change of >2 were plotted using ggplot2. Gene Ontology analysis was performed using the goseq package using hg19 build.^5^ The semantic similarity-based scatterplot of GO terms was created using REVIGO (http://revigo.irb.hr).^6^ The heatmap shown in Supplementary Figure 1 was performed using the Biojupies platform.^7^ Sequencing data of control cell lines were obtained from the Gene Expression Omnibus public repository (NIH). Prior to all analysis, the raw gene counts were normalized using the logCPM method by dividing each column by the total sum of its counts, multiply by a million and followed by the application of a log10-transform. Genes with the most variable expression were selected and transformed using the Z-score. Principal Component Analysis was generated using the Plotly's Python graphing library and sklearn Python module. Heatmap was created using Clustergrammer.^8^

**Dimensionality reduction of RNAseq datasets using UMAP**

Raw counts from different RNA-seq datasets were normalized (divided by the total count, multiplied by 1 million and natural-log transformed using log1p) and center-scaled. From the scaled data, the most variable genes (3,000) across datasets were used to conduct principal component analysis and the top principal components (17) were selected for dimensionality reduction and 2-dimention representation using the UMAP (Uniform Manifold Approximation and Projection) and DimPlot functions from the R Package Seurat.^9–11^

**Data availability**

The RNA sequencing data discussed in this publication have been deposited in the NCBI Gene Expression Omnibus under accession number GEO: GSE152390.

(<https://www.ncbi.nlm.nih.gov/geo/query/acc.cgi?acc=GSE152390>)

### Supplementary Figure Legends

**Supplementary Fig. S1. Pluripotent Stem Cells’ characterization. A)** Representative morphology of one of the induce pluripotent stem cell (iPSC) populations (phase contrast, scale bar 1000 µM left, 200 µM right); **B)** No difference in the expression of pluripotency genes was noted between iPSC and embryonic stem cells (ESC; RT-qPCR, expressed as log_10_ mean fold change ± SEM; n=3). **C)** Expression of pluripotency markers in iPSC compared to primary human fibroblasts (immunofluorescence, representative image, DAPI nuclear staining in included images, scale bar 200 µM). **D)** Expression of pluripotency markers was highly homogeneous in used iPSC populations (flow cytometry, representative experiment). **E)** Gene expression analysis showing that the iPSC and ESC populations used for this paper were undistinguishable from previously published iPSC and ESC populations, while clearly very different from PBMC, embryonic fibroblasts and CD34+ cells (heatmap showing the expression of the top 2500 most variable genes across samples; RNA-Seq, unsupervised clustering).

**Supplementary Fig. S2. A-B) Effect of knock-our serum replacement (KOSR) on cell viability and quality of differentiation:** supplementation with 1% KOSR significantly increased survival over the first 5 days of differentiation (A; n=3) with both our new differentiation protocol (ProtB) and a protocol not acting on Wnt and TGFβ signaling pathways beyond the definitive endoderm stage (ProtA), with no significant effect on the expression of endoderm-related genes (B; n=3). **C-D)** **No difference was noted between iPSC and ESC** in the results obtained at the stages of definitive endoderm (C) and liver bud hepatoblasts (D; RT-qPCR, expressed as log_10_ mean fold change ± SEM; n=3). **E) The differentiation protocol proved to be robust**, with little intra- and inter-population or inter-operator variability (32 differentiations of 6 PSC populations, 3 different operators; RT-qPCR, expressed as log_10_ mean fold change; *p*=n.s.). **F)** **No significant difference was noted between HLC obtained from iPSC and ESC** (RT-qPCR, expressed as log_10_ mean fold change ± SEM; n=3).

**Supplementary Fig. S3. Schematic representation of the differentiation protocol used as comparison** (protocol not acting on Wnt/β-catenin and TGFβ signaling pathways beyond the definitive endoderm stage, see supplementary methods for details).

**Supplementary Fig. S4. Comparison between HLC obtained with our new protocol and those obtained with a protocol not acting on Wnt and TGFβ signaling pathways beyond the definitive endoderm stage (HLC-2).** A) Heatmap showing the expression of the top 500 most variable genes across samples (RNA-Seq, 2 samples per group, unsupervised clustering). B) Volcano plot showing the most variable genes in our HLC compared to HLC-2 (n=2). C) Representative sample of liver-enriched genes overexpressed in HLC obtained with the new protocol compared with HLC-2 (differential gene expression; >3-fold change, p-adjusted <0.05; n=2). D) Semantic similarity-based scatterplot of Gene Ontology terms showing families of genes that are overexpressed in HLC compared to HLC-2 (REVIGO tree map, n=2, p-adjusted <0.05).

**Supplementary table 1. TaqMan gene expression assays used for RT-qPCR.**

**Supplementary table 2. Primary antibodies used for immunofluorescence.**
