## Supplementary figures and images for "Leveraging complex interactions between signaling pathways involved in liver development to robustly improve the maturity and yield of pluripotent stem cell-derived hepatocytes"

### Supplementary Fig. S1

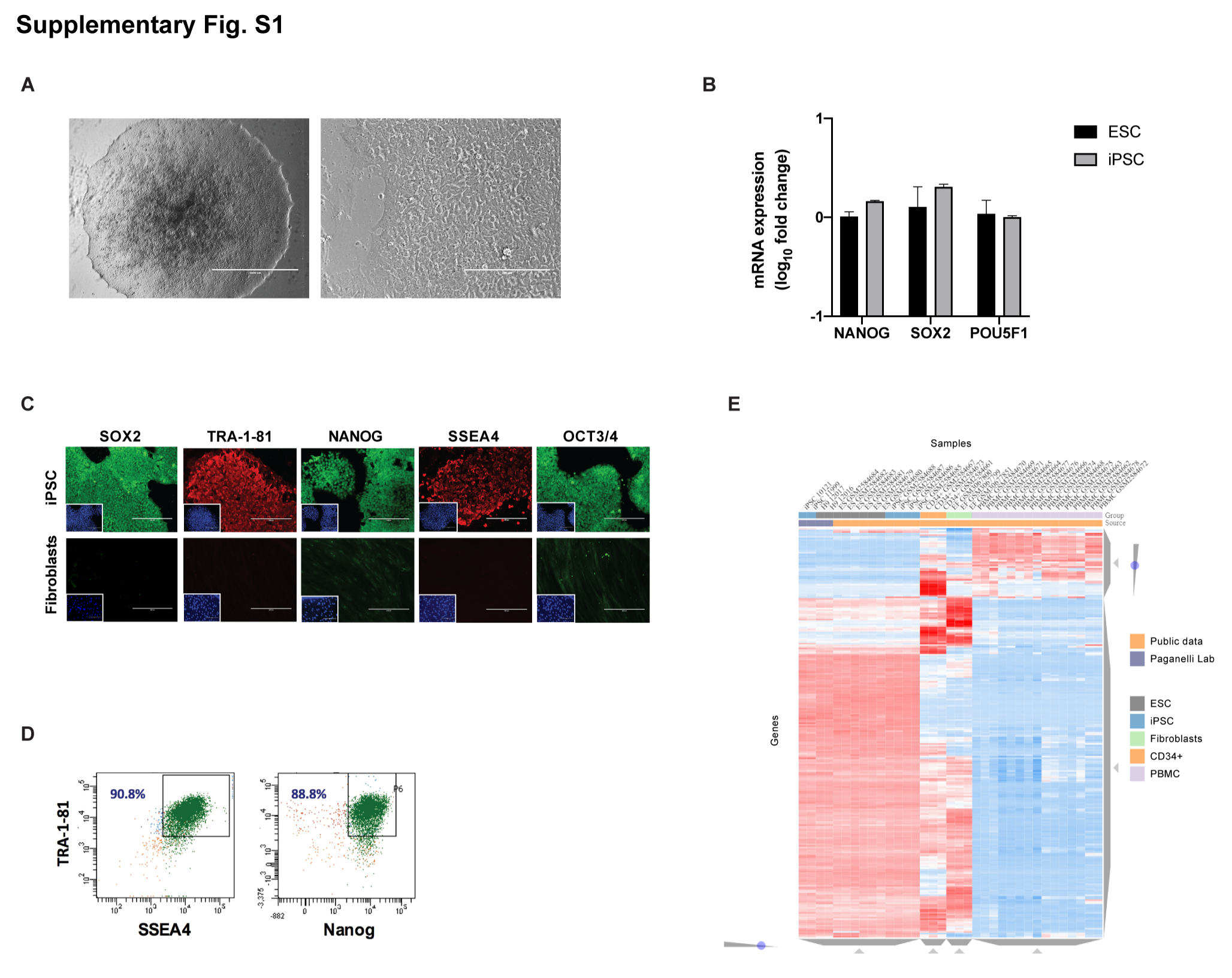

### Supplementary Fig. S2

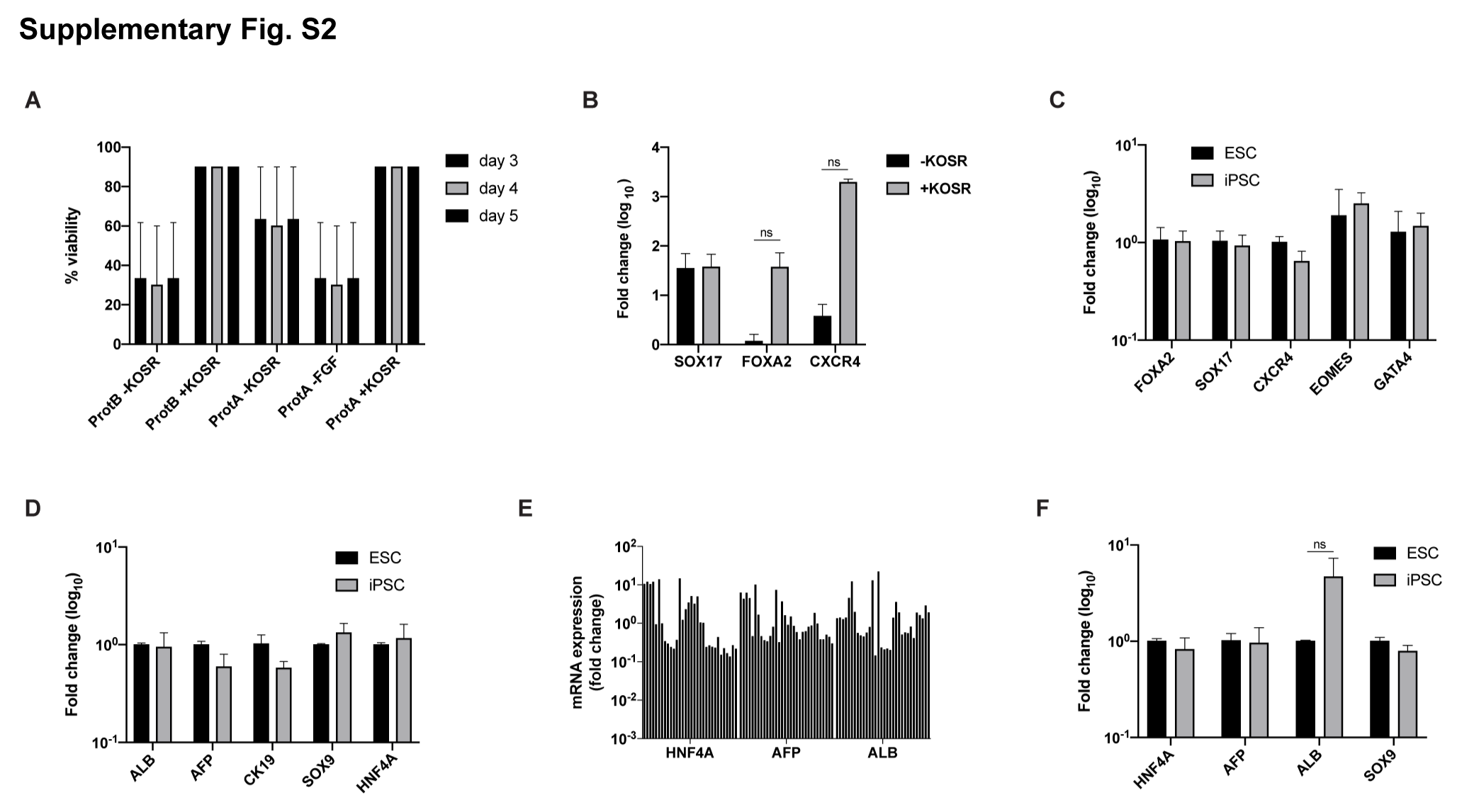

### Supplementary Fig. S3

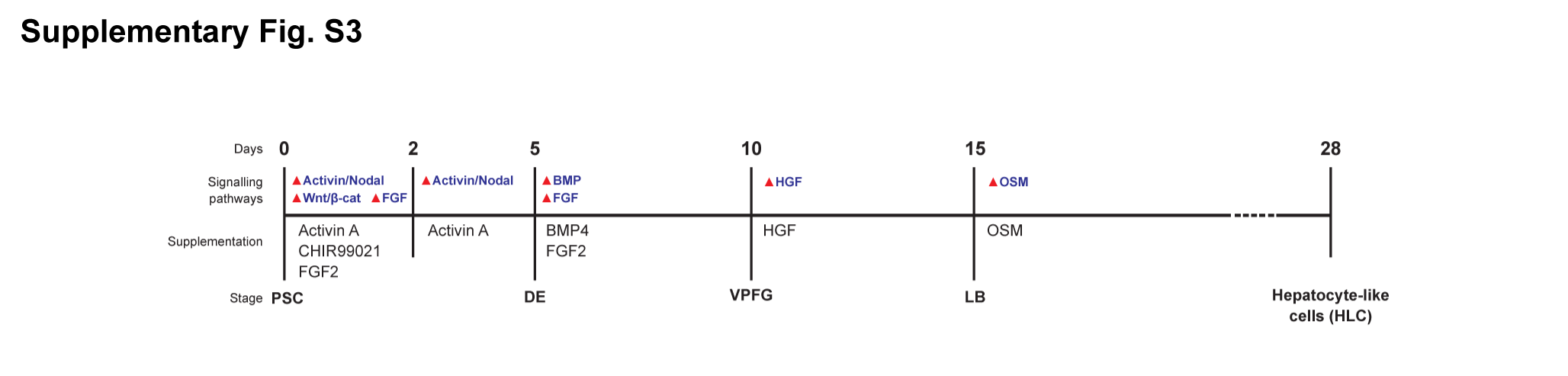

### Supplementary Fig. S4

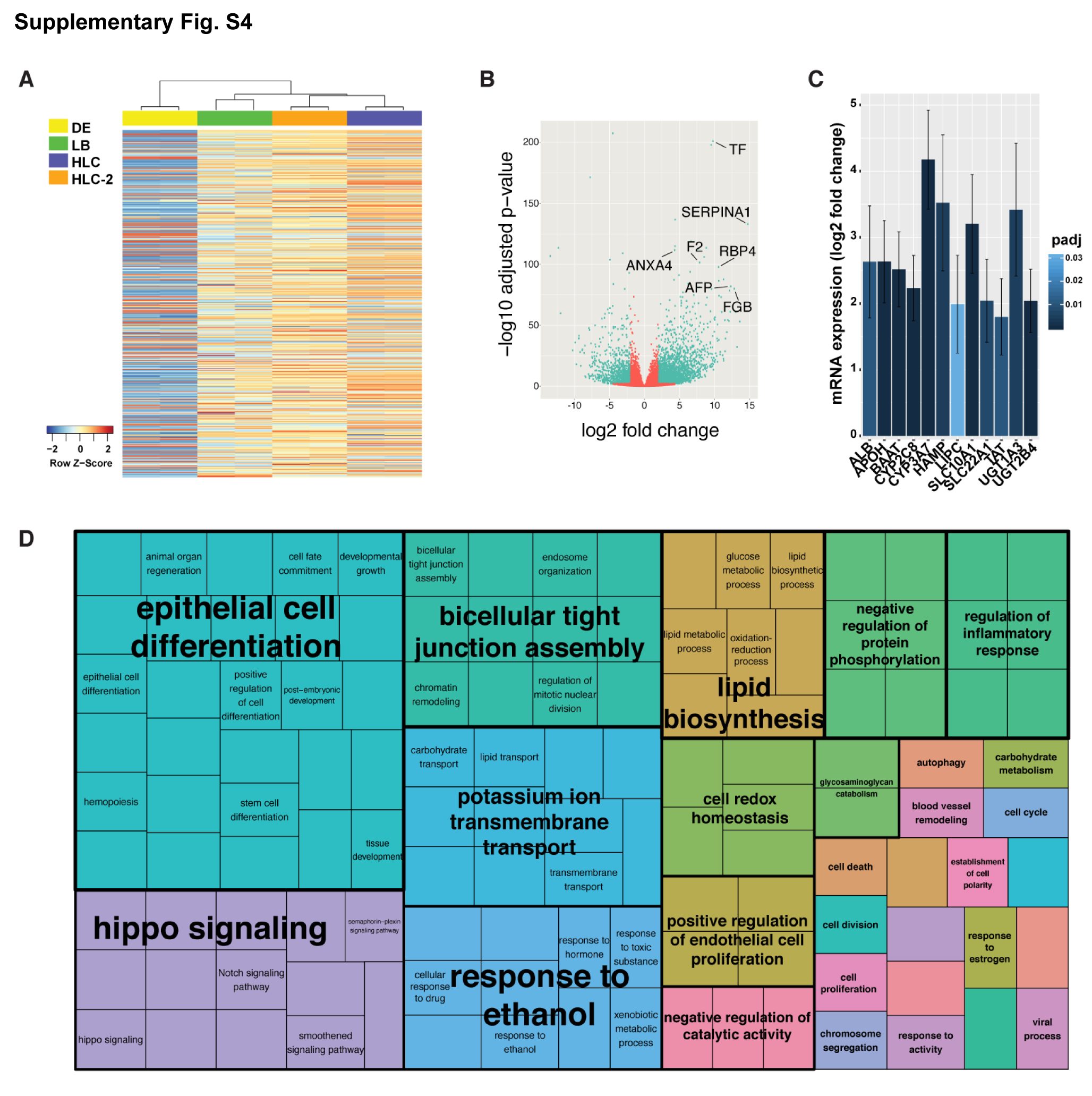
